# Mining the potential of *VRS1-5* gene to raise barley grain yield

**DOI:** 10.1101/2020.09.22.307876

**Authors:** Liping Shen, Yangyang Liu, Zhiwen Sun, Ziying Wang, Lili Zhang, Yu Cai, Yuannian Jiao, He Wu, Kuocheng Shen, Ping Yang, Zifeng Guo

## Abstract

*VRS1-5* genes determine spike row types during the early stages of spike development in barley (*Hordeum vulgare*), yet their functions for the determination of grain yield during the late stages of spike development are largely unknown. To assess the role of *VRS1-5* genes in determining grain yield components, we sequenced *VRS1-5* genes from 894 worldwide barley accessions and measured 19 spike morphology traits in four environments. Single nucleotide polymorphism SNP markers and gene marker-based haplotypes for *VRS1-5* displayed close associations with spike morphology traits. We further developed a spatiote-temporal transcriptome atlas (255 samples) at 17 stages and five positions along the spike, that linked spike morphology to spikelet development and expression patterns of *VRS1-5* genes. Phenotypic measurements demonstrated that mutations in *VRS1-5* suppress the initiation of spikelet primordia and, trigger spikelet abortion by increasing cytokinin content and improving sensitivity of spikelet primordia to cytokinin. Our integrated results illustrate how breeding can globally alter spike morphology through diversity at the *VRS1-5* genes, which show great potential in increasing barley grain yield.

## Introduction

Inflorescence displays variable architectures in the grass family. In temperate cereal crops, the grain-bearing inflorescence is called a spike that consist of several basic “spikelet” units. A spikelet is itself composed of one or more florets, which include the floral organs, e.g. lemma, palea, lodicules, stamens and carpels. In barley (*Hordeum vulgare*), each rachis node is composed of triplets (each with one central spikelet and two lateral spikelets). Depending on the fertility of the two lateral spikelets, each rachis node will bear either two-rowed or six-rowed spikes. In wild barley (*Hordeum vulgare ssp. spantaneum*), the progenitor of modern-day cultivated barley, one rachis carries one fertile central spikelet and two sterile lateral spikelets, forming a two-rowed spike. In modern six-rowed barley cultivars, all three spikelets are fully fertile and able to develop into grains at each rachis node. Six-rowed barley arose during barley domestication through the selection of spontaneous recessive variants with the six-rowed phenotype about 8,000-12,000 years ago (1, 2). To date, five genes have been shown to independently regulate spike transition from two-rowed to six-rowed spikes: *SIX-ROWED SPIKE 1* (*VRS1*), *VRS2, VRS3, VRS4* and *VRS5* (3–8). The determination of barley row type is accomplished during the awn primordium (AP) stage (the stage at which the maximum number of spikelet primordia is present, when spike length is ∼ 0.5–1.0 cm). However, the role of *VRS1-5* in determining grain number and spike development after the AP stage is not well illustrated.

The survival rate of spikelet primordia per spike after the AP stage plays a critical role in determining grain yield in barley, as a big proportion of spikelet primordia within an individual spike will abort (9–12). before grain setting takes place at physiological maturity. After reaching the maximum number of spikelet primordia at the AP stage, spikelet primordia start to disintegrate. Aborted spikelet primordia almost disappear by the time the inflorescence reaches the tipping (TP) or heading (HD) stages. In particular, the degeneration of apical spikelet primordia causes the largest loss of grain number potential. In addition, after the degeneration of spikelet primordia, the spikelets with differentiated organs (e.g. anthers, ovary) at one or two additional rachis nodes (one spikelet at each rachis node in two-rowed barley; three spikelets at each rachis node in six-rowed barley) will degenerate. These later degenerated spikelets will not set grains, although the empty spikelets remain visible until physiology maturity.

Understanding the molecular mechanisms underlying spikelet development and degeneration is vital for improving grain yield in barley. To address this, we sequenced *VRS1-5* in 894 worldwide barley accessions to explore their coding diversity. We further developed a spatio-temporal transcriptome atlas across 17 spikelet developmental and degeneration stages and at five rachis node positions to show the expression patterns of *VRS1-5*. Moreover, we determined that *vrs1-5* mutants affect cytokinin contents, and thus spike architecture, on spikelet primordia nodes.

## Results

Different barley accessions show variable spike morphologies (Fig. 1). Barley breeding selected obvious modifications in spike morphology to improve grain yield and adaptation to the local environment. In this study, we investigated the potential for *VRS1-5* genes to increase barley grain yield. To assess the connections between spike morphology and *VRS1-5*, we measured 19 spike morphology and sequenced the partial coding regions (800-1000bp) for *VRS1-5* in 894 worldwide barley accessions from 35 countries (Table S1).

**Fig. 1.**
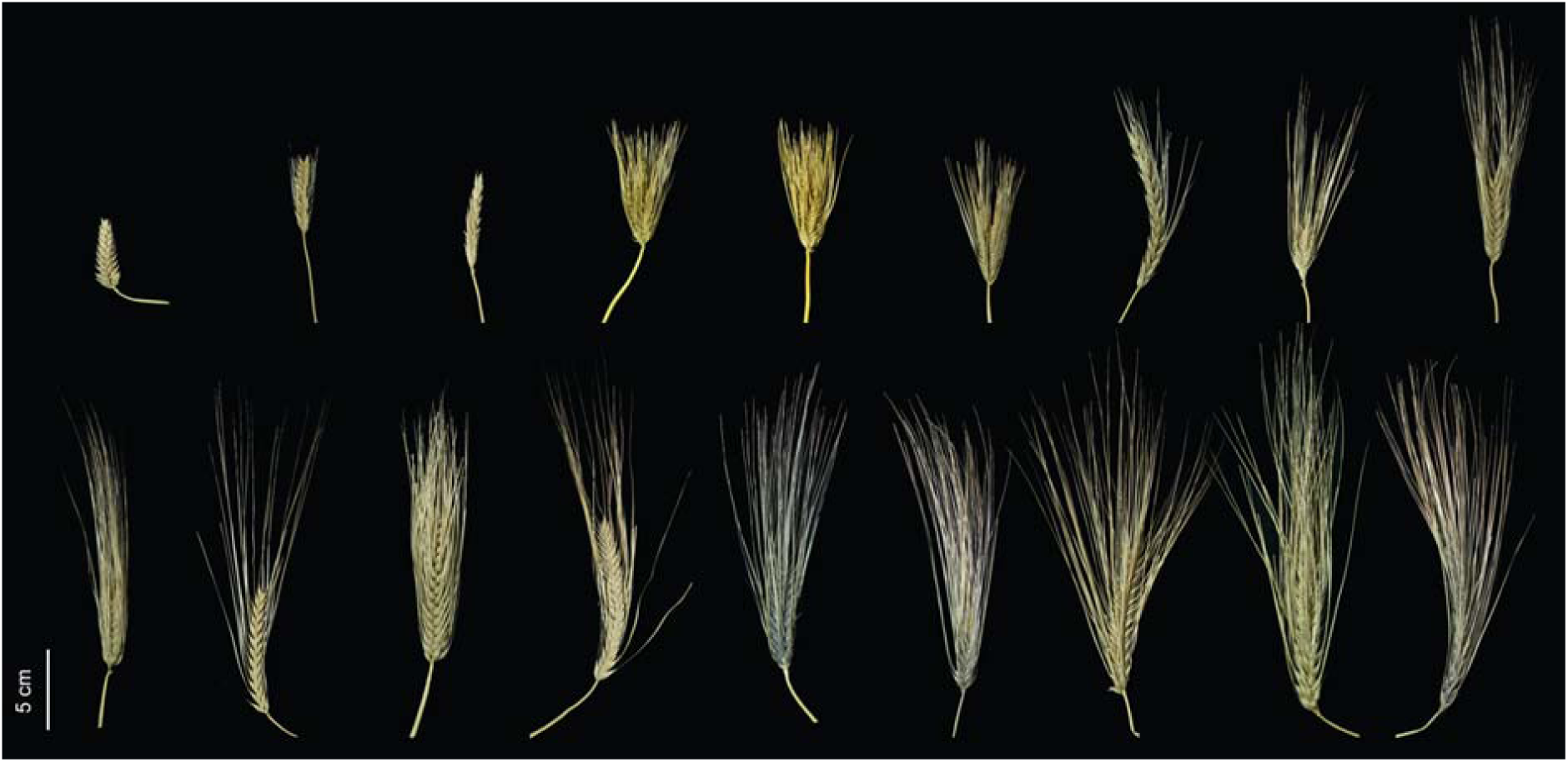
Variation in spike morphology across barley accessions illustrates the potential of untapped genetic regulation for the improvement of grain yield though manipulation of spike morphology traits.

### Associations between single nucleotide polymorphism (SNP) markers, gene marker-based haplotypes at *VRS1-5* and spike morphology in barley

Since genetic diversity is a reflection of nucleotide diversity and haplotype diversity, we computed the haplotypes and single nucleotide polymorphism (SNPs) for *VRS1-5*. Based on their coding regions in 894 worldwide barley accessions, we detected SNP markers (with a minor allele frequency, MAF > 0.05) for each *VRS*: 25 for *VRS1*, 33 for *VRS2*, 20 for *VRS3*, 26 for *VRS4*, and 22 for *VRS5*. We identified 29 haplotypes (with each haplotype including more than 10 barley accessions) in our barley VRS sequence collection, consisting of 4 haplotypes for *VRS1*, 7 for *VRS2*, 2 for *VRS3*, 10 for *VRS4* and 6 for *VRS5* (Fig. 2a,2b,2c, Fig.S1, Fig. S2, Table S2-6). We detected most haplotypes in accessions originating from all 35 countries represented in our collection. In addition, for spike morphology traits, we identified 52 SNP marker-trait associations (MTAs, P<0.005) for *VRS1*, 297 for *VRS2*, 60 for *VRS3*, 194 for *VRS4*, and 390 for *VRS5* across the four environments tested here (Table S7).

**Fig. 2.**
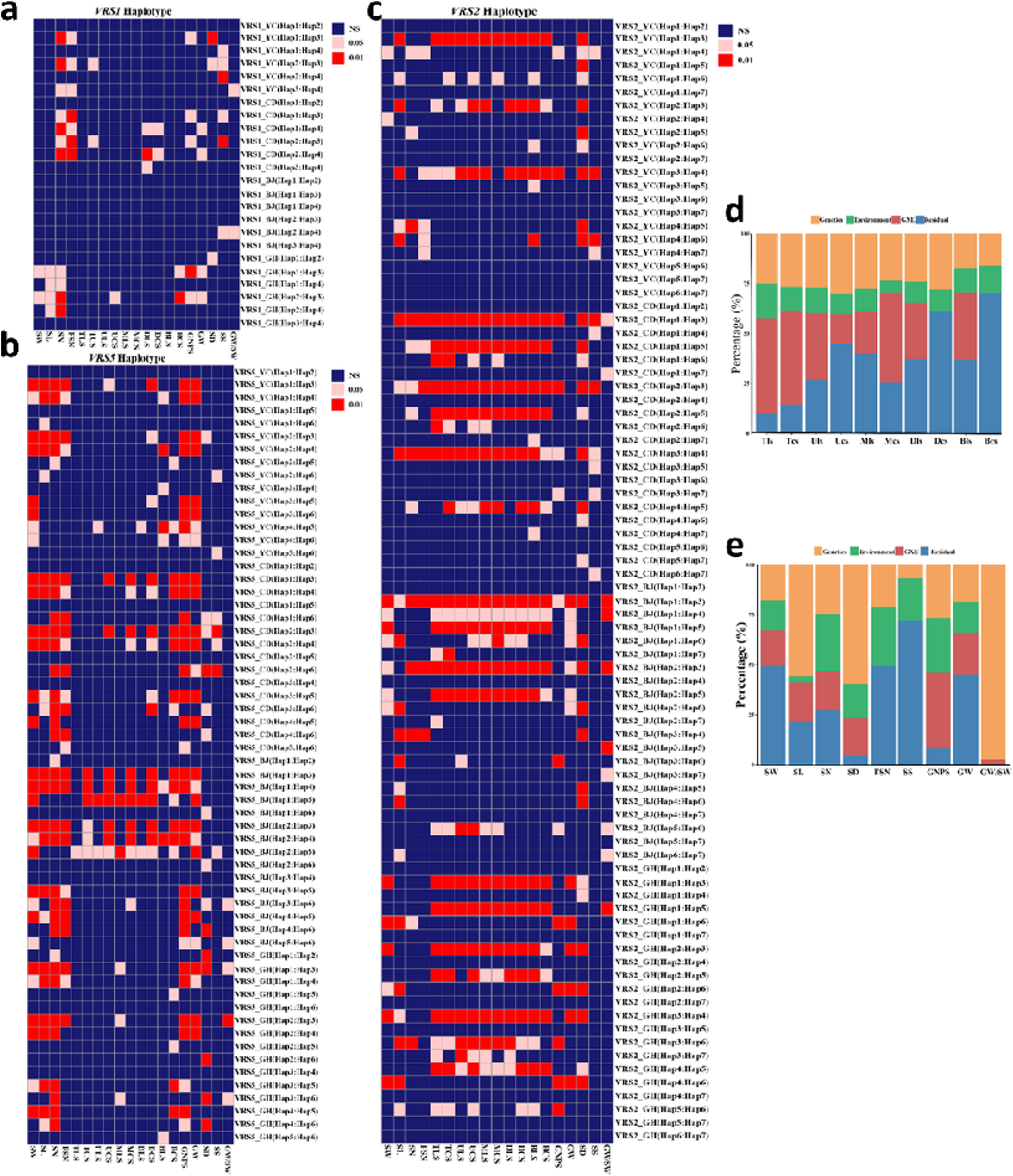
The effects of variation at the *VRS1-5* genes on spike morphology traits. (a-c) Differences for 19 spike morphology traits between haplotypes for *VRS1* (a), *VRS5* (b), and *VRS2* (c), respectively. (d–e) Analysis of variance (ANOVA) for the 19 spike morphology traits. SW, spike weight; SL, spike length; SN, spikelet/rachis node number; FSN, fertile spikelet/rachis node number; TLS, top lateral spikelet weight; TCS, top central spikelet weight; ULS, lateral spikelet weight at the position of 25% spike length from top; ULS, central spikelet weight at the position of 25% spike length from top; MLS, lateral spikelet in the middle position of spike; MCS, central spikelet in the middle position of spike; DLS, lateral spikelet weight at the position of 75% spike length from top; DLS, central spikelet weight at the position of 75% spike length from top; BLS, lateral spikelet in the bottom of spike; BCS, central spikelet in the bottom of spike; GNPS, grain number per spike; GW, grain weight per spike; SD, spikelet density, the ratio between spikelet/rachis node number and spike length; SS, spikelet survival, the ratio between spikelet/rachis node number and fertile spikelet/rachis node number; GW/SW, the ratio between grain weight and spike weight.

We identified four haplotypes for *VRS1*, each grouping 589 accessions (Hap1), 230 accessions (Hap2), 72 accessions (Hap3) and 18 accessions (Hap4) out of the 894 cultivars evaluated here. Compared to that of Hap1 and Hap2, the presence of the Hap3 haplotype significantly increased rachis node number and fertile rachis node number in accessions grown in the south-eastern, and south-western locations and in the greenhouse (Table S8). Similarly, Hap4 increased rachis node number and fertile rachis node number in both the south-western location and in the greenhouse relative to the Hap1 and Hap2 haplotypes (Fig. 2a). We did not observe any significant differences in lateral or central spikelet dry weight between haplotypes.

We detected 7 haplotypes for *VRS2*, each haplotype consisting of388 accessions (Hap1), 171 accessions (Hap2), 95 accessions (Hap3), 59 accessions (Hap4), 30 accessions (Hap5), 28 accessions (Hap6) and 11 accessions (Hap7). We observed clear differences for all 19 spike morphology traits across the seven *VRS2* haplotypes of *VRS2*, with additional contribution from the environment, For lateral and central spikelet dry weight, we noticed clear differences between five haplotype combinations in the northern location: Hap2-Hap7, Hap3-Hap4, Hap3-Hap7, Hap4-Hap5, Hap5-Hap7; We also observed differences between three haplotype combinations in the south-eastern location: Hap2-Hap3, Hap3-Hap4, Hap3-Hap7; For the two remaining locations, we detected five haplotype combinations in the south-western location: Hap2-Hap3, Hap3-Hap4, Hap3-Hap7, Hap4-Hap5, Hap5-Hap7; and six haplotype combinations when plants were grown in the greenhouse: Hap2-Hap3, Hap2-Hap5, Hap3-Hap4, Hap3-Hap7, Hap3-Hap9, Hap4-Hap5, Hap5-Hap7 (Fig. 2c, Table S8).

For *VRS3*, we identified two haplotypes in 632 (Hap1) and 254 (Hap2) accessions. We did not detect significant differences between the 19 spike morphology traits between these two haplotypes, except one trait: spikelet density (Fig. S1, Table S8). However, MTAs showed connections between *VRS3* and spike morphology (Table S7).

For *VRS4*, we detected ten haplotypes, grouping together 270 (Hap1), 169 (Hap2), 127 (Hap3), 119 (Hap4), 58 (Hap5), 43 (Hap6), 31 (Hap7), 19 (Hap8), 12 (Hap9) and 12 (Hap10) accessions. All 19 spike morphology traits displayed clear differences across the 10 haplotype combinations (Fig. S2, Table S8). These effects of *VRS4* haplotypes on spike morphology traits were identical to those seen with *VRS2*: haplotype combinations displayed different effects in the four environments (Fig. S2, Table S8).

Finally, we detected six haplotypes for *VRS5*, each consisting of 571 (Hap1), 157 (Hap2), 71 (Hap3), 63 (Hap4), 53 (Hap5) and 20 accessions (Hap6). *VRS5* haplotypes showed obvious effects on nine spike morphology traits, but not on lateral or central spikelet dry weight at five positions, especially with plants grown in the greenhouse and in the southwestern location (Fig. 2b, Table S8).

The results of variance components analyses reveal that: environmental effects (σ^2^ _E_ and σ^2^ _GXE_) contributed largely to the lateral spikelet (65%) and central spikelet (59%) at top of spike as well as grain number per spike (65%) (Fig. 2d, Table S9); genetic factors (σ^2^ _G_) display great contribution to spike length (56%), rachis node density (60%) and the ratio between grain weight and spike dry weight (98%) (Fig. 2e, Table S9).

In summary, the haplotypes of *VRS1, VRS2, VRS4*, and *VRS5* displayed clear effects on all spike morphology traits, but we observed no significant differences for lateral or central spikelet dry weight among the *VRS1* and *VRS3* haplotypes. We also did not dectect obvious effects from the *VRS3* haplotypes on spike morphology, which may be attributed to the small number of *VRS3* haplotypes in this study. The different effects of haplotypes, the different MTA among the four environments as well as the results of variance components analyses indicate the contributions of environments and interactions between *VRS1-5* and environments for the spike morphology traits.

### Spatiotemporal dissection of spike transcriptome links spikelet development and *VRS1-5* expression patterns

Spikelet primordia abortion within individual spike is responsible for a 30–50% loss in grain yield potential. Before grain setting at physiological maturity, spikelet primordia suffer a large loss of spikelet primordia (Fig. 3a) during the development and abortion process (Fig. 3b).

**Fig. 3.**
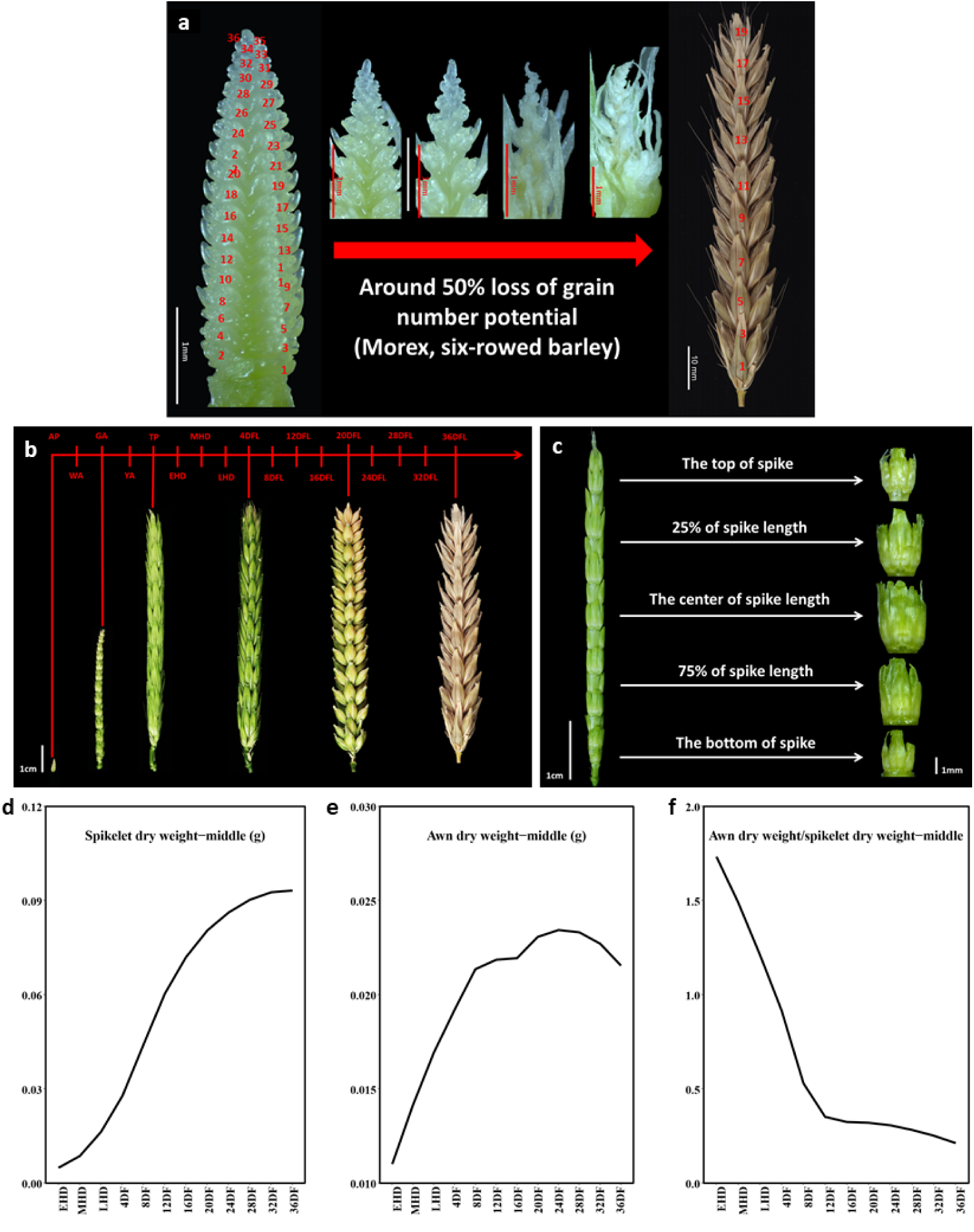
Spatiotemporal dissection of spike development in barley. (a) Floret/floret primordium abortion (the top of the spike) across the 17 spike developmental stages. The first row displays floret primordium abortion, which means these floret primordia abort before they develop into florets. In addition, 1–2 florets may abort even though they already have anther, ovary and other floret structures after the tipping stages. (b) Temporal dissection-of spikelet development into 17 stages: awn primordium (AP) stage, white anther (WA) stage, green anther (GA) stage, yellow anther (YA) stage, tipping stage (TP), early heading stage (EHD), middle heading (MHD) stage, late heading (LHD) stage, 4days after late heading stage (4DFL), 8 DFL, 12DFL, 16DFL, 20DFL, 24DFL, 28DFL, 32DFL, 36DFL. The samples for transcriptome analysis were collected at each of these17 stages. Spikes at the AP, GA, TP, 4DFL, 20DFL and 36DFL (physiological maturity stage) stages are shown to illustrate the dynamic change in spike morphology at these 17 stages. (c) Spatial dissection of the five spikelet positions within individual spikes: spikelets at the top of the spike, spikelets along the top 30% of the spike, spikelets at the center of the spike, spikelet along the bottom 30% of the spike, spikelets at the bottom of the spike. (d-f) Trends for weight of three spikelets (Morex is a six-rowed barley variety,with three spikelets per node) at each node, weight of awn at each node, and the ratio between awn weight and spikelet weight.

To monitor the dynamics in gene expression during the development and abortion process of spikelet primordia, we undertook a deep-sequencing of the transcriptome (RNA-Seq) approach across 17 developmental stages: (AP) stage, white anther (WA) stage, green anther (GA) stage, yellow anther (YA) stage, tipping (TP) stage, early heading (EHD) stage, middle heading (MHD) stage, late heading stage (LHD, more that 90% of spike visible), 5 days after late heading (5 DFL) stage, 10DFL, 15 DFL, 20DFL, 25DFL, 30DFL, 35DFL (Fig. 3b). To determine gene expression profiles across the unbalanced developmental and abortion gradient of spikelet within individual spikes, we also performed an RNA-Seq analysis of spikelet at five positions along individual spike: spikelets at the top of the spike, spikelets at the top 30% of the spike, spikelets at the center of the spike, spikelet at the bottom 30% of the spike, spikelets at the bottom of the spike (Fig. 3c). This spatiotemporal gene expression dataset of spikelets at 17 stages and five positions allowed us to determine the dynamics and differences in gene expression profiles according to stages of spikelet development and degeneration process.

#### (1) Trends in spike architecture traits and gene expression patterns

We divided spike development into 17 stages (Fig. 3b). We monitored plant growth at 17 specific stages and 5 spikelet positions by measuring spike morphology traits: awn dry weight DW, lateral spikelet DW, central spikelet DW and total spikelet DW at 12 stages and five spikelet positions. For all spikelet positions, we noticed fast growth and a large increase in lateral spikelet DW, central spikelet DW and total spikelet DW between the TP and 24DF stage, which corresponds to the grain filling process. Awn DW increased quickly between the GA and 24DF stages, which therefore did not quite follow the same trend as spikelet DW. Notably, the ratios between awn DW and spikelet DW strongly decreased from the GA to the 8-DF stage. All measured traits displayed variable trends among the 17 developmental stages, indicating that different traits reached their peak values at specific and distinct stages (Fig. 3d, 3e, 3f, Fig. S3, S4, S5, S6, S7). Moreover, temporal trends of five spikelet positions are identical (Fig. S3, S4, S5, S6, S7).

To further explore the functional transitions during spike development, we clustered all expressed genes as five modules according to their expression patterns, and then performed functional categori for each module (Fig. 4a, 4b; Fig. S8, S9, S10, S11). For the apical spikelet position of the spike, the AP-WA module contained a set of genes related to floral organ initiation and development (Fig. 4b), which is consistent with what is happening in barley between the AP and WA stage: floral organs (e.g. anther, ovary) initiate and differentiate; We also determined that GA-YA cluster was enriched in genes with categories related to pollen and ovary development (Fig. 4b), which agrees with pollination taking place around the YA stage and thus require both male and female gametes. From the TP to the HD stage, fast grain filling occurs, which necessitates a large and constant influx of carbohydrate, while the spike is losing water (Fig. 4b).in agreement, we observed an enrichment in such categories as “carbohydrate metabolic process “and “response to water deprivation” in this phase. From 4 DFL to DFL36, the grain is hardening and filling (Fig. 4b), a phase during which the grain is sensitive to the stresses (e.g. heat, drought) and highly responsive to phytohormones. In addition, we noticed that the functional categories for modules associated with each of the five spikelet positions were identical (Fig. S8, S9, S10, S11).

**Fig. 4.**
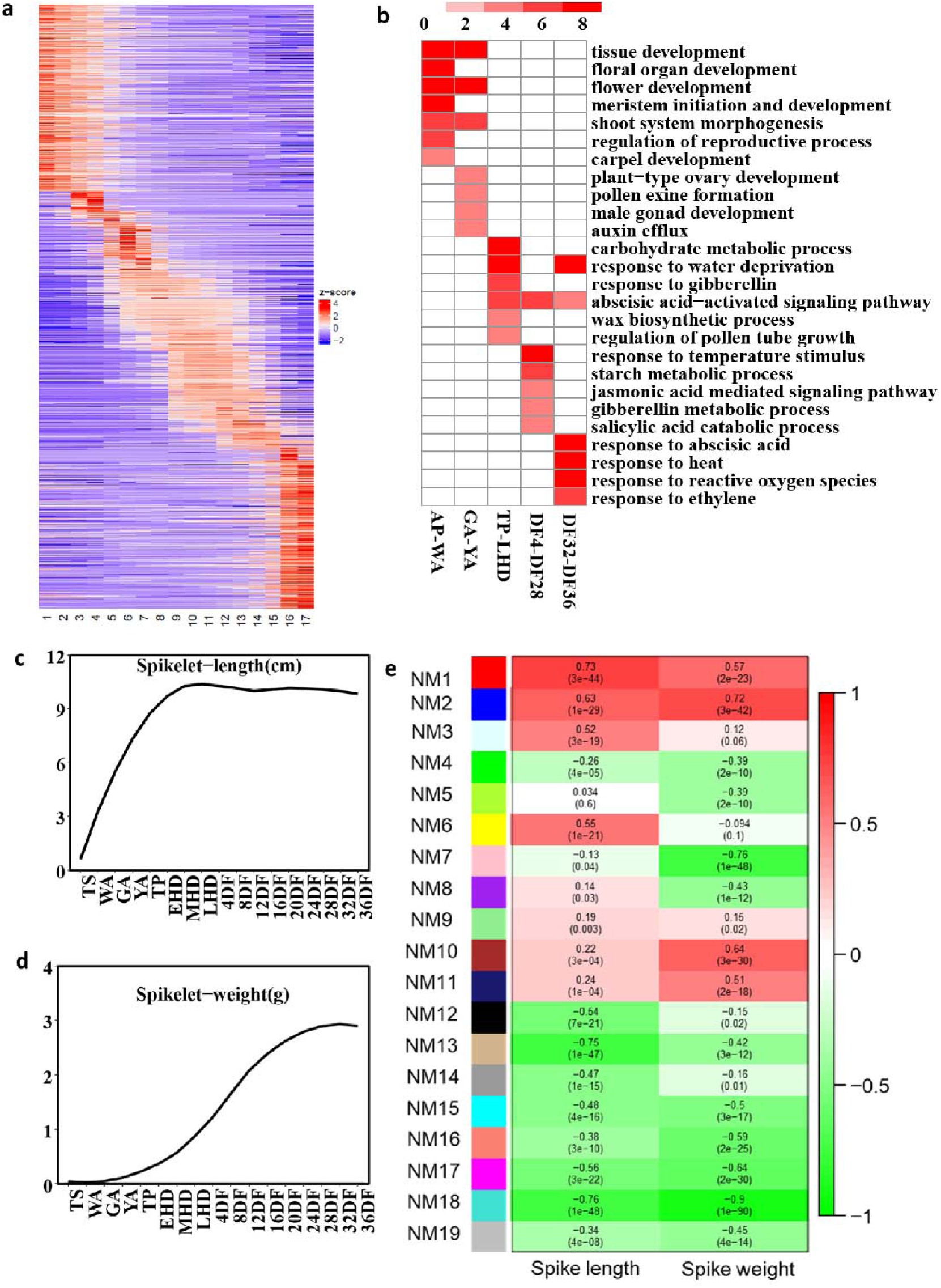
Connections between gene functions and spikelet development. (a) Expression patterns of genes at the apical spikelet position of spike across the 17 developmental stages. Red: induced gene; blue: repressed gene. (b) Functional categories enriches in different groups based on the five modules of 17 spike developmental stages at apical spikelet position of spike. Red: logarithm (log_2_()) of relevant gene number. (c-d) Trends in spike length (c) and spike weight (d) across the 17 developmental stages. (e) Associations between spike morphology traits and gene modules. The colors indicate the correlation values.

#### (2) Network topology between spike architecture and gene module

Spike length rapidly increased from AP to EHD, after which spike stopped elongating until 36DF (Fig. 4c). Spike DW rose slowly from the AP to the MHD stage, then increased rapidly from the MHD to the 24 DF stage, and stayed constant from 24 DF to 28DF (Fig. 4d). To categorize the transcriptionally-regulated biological processes in spikelet degeneration and development, we performed a weighted gene co-expression network analysis (WGCNA) of all 30060 expressed genes to identify representative network modules (NMs) that were highly associated with spike DW and length (Fig. 4e), resulting in 19 NMs.

We assigned putative biological functions to the 19 NMs by performing Gene Ontology (GO) enrichment analyses. GO terms were highly consistent with functional transitions (Fig. 4b). Module NM2 contained genes associated with starch metabolism, response to heat, hormone−mediated signalling pathway, and cellular response to endogenous stimulus. Module NM4 was enriched in genes involved in tissue development, regulation of programmed cell death, regulation of growth rate, pollination and, carpel development. Module NM8 was enriched in genes related to regulation of reproductive process, regulation of pollen tube growth, pollen tube development and developmental maturation (Table S10). The GO terms enriched for NM16 included all the functional categories associated with NM2, NM4, and NM8 (Table S10). In addition, NM6 was enriched in genes with roles in response to water deprivation, response to fructose, response to water, response to carbohydrate, and drought recovery (Table S10).

#### (3) *VRS 1-5* expression patterns

To investigate the potential connections between *VRS1-5* and spikelet development and degeneration, we determined their gene expression patterns across all 17 developmental stages (Fig. 5, Fig. S12, S13, S14, S15). The expression of *VRS2* and *VRS5* decreased dramatically from AP/WA to YA/TP, suggesting that these two genes play important roles before pollination. *VRS1* had the highest expression level relative tothe other four VRS genes, suggesting that *VRS1* may be important throughout spike development. However, we did observe two peaks in *VRS1* expression at the YA and EHD stages, indicating that *VRS1* may take on a critical role during the early late pollination phase and the early grain filling phase. *VRS3* and *VRS4* followed a very similar pattern across all 17 stages and five spikelet positions; they both rose starting at the AP stage and reached their peak at the TP stage. Thereafter, their expression decreased markedly, suggesting that *VRS3* and *VRS4* may contribute to the regulation of early grain filling phased.

**Fig. 5.**
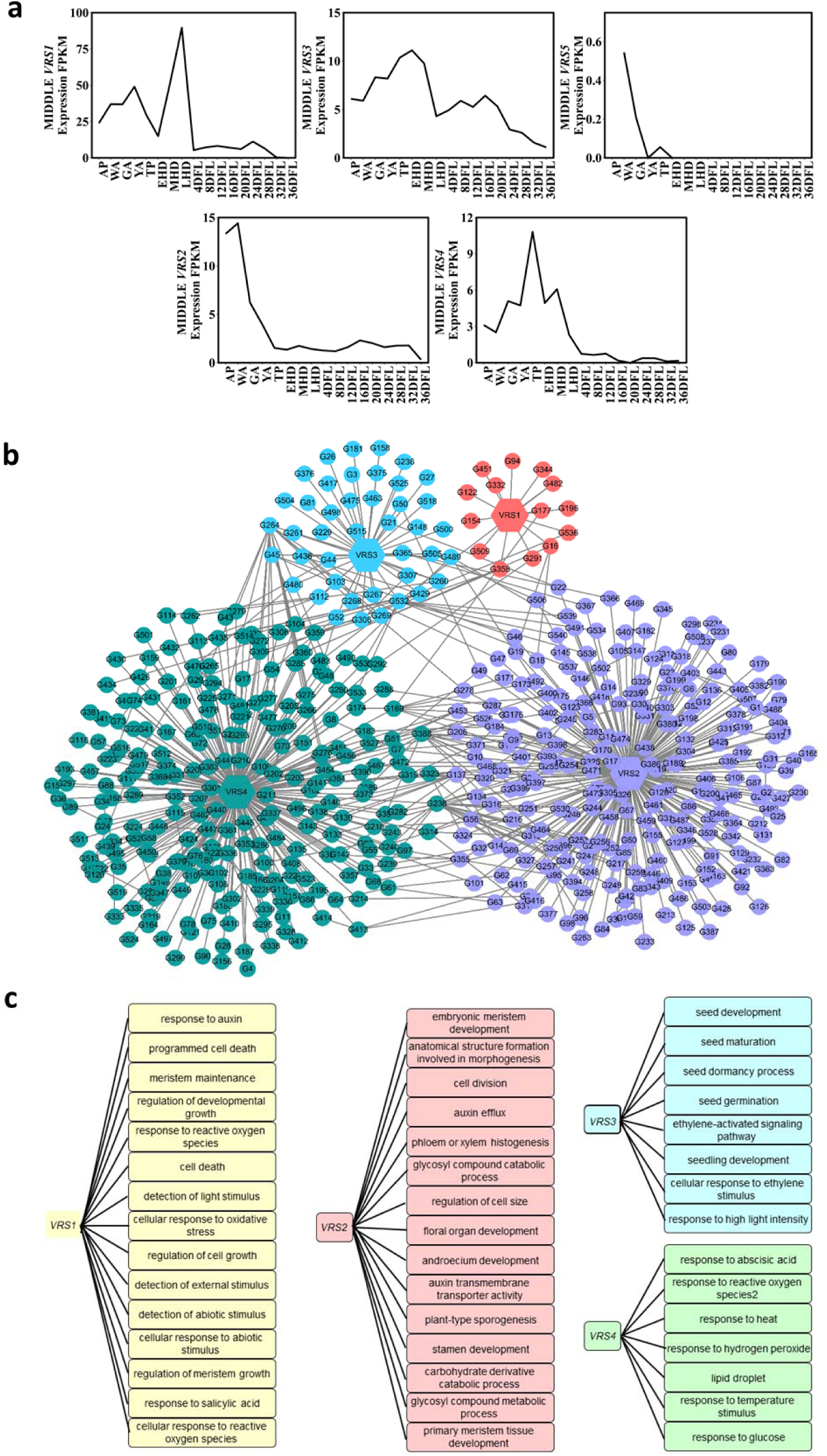
The associated genes of *VRS1-5* according to the spatiotemporal transcriptome analysis. (a) Expression patterns of *VRS1-5* genes at central spikelets of spike across the 17 spikelet developmental stages in barley. (b) The network of *VRS1*-4 and their associated genes in the same co-expression network analysis (WGCNA) module, *VRS5* was not included in any WGCNA module due to its low expression. (c) the consistent Gene Ontology (GO) terms of *VRS1*-4 associated genes with functional transitions along spike developmental process (Fig. 4b) as well as some functional categories in the WGCNA modules (NM2, NM4, NM6, NM8, NM16) (Fig. 4e, Table S10).The information for the 17 stages can be found in Fig. 3: awn primordium (AP) stage, white anther (WA) stage, green anther (GA) stage, yellow anther (YA) stage, tipping (TP) stage, early heading (EHD) stage, middle heading (MHD) stage, late heading (LHD) stage, 4days after late heading stage (DFL), 8 DFL, 12DFL, 16DFL, 20DFL, 24DFL, 28DFL, 32DFL, 36DFL.

We determined the associations between *VRS1*-4 and the genes in the same WGCNA module (*VRS3* in WGCNA module NM10, *VRS4* and *VRS2* in WGCNA module NM18, *VRS1* in WGCNA module NM7, Fig. 4e), *VRS5* was not included in any WGCNA module due to its low expression. The network between *VRS1*-4 and their 540 associated genes suggested the connections between *VRS1*-4 as well as their associations with genes of viable functions (Fig. 5b, Table S11). To further assess the functional categories of the 540 genes which are associated with *VRS1*-4, we further conducted Gene Ontology (GO) enrichment analyses. The GO terms of *VRS1*-4 associated genes (Fig. 5c) cover identical functional categories that were identified in functional transitions along spike developmental process (Fig. 4b) as well as in the WGCNA modules (NM2, NM4, NM6, NM8, NM16) (Fig. 4e, Table S10). For example, *VRS1* associated genes was enriched for terms including cell death, detection of external stimulus, response to auxin, response to salicylic acid, meristem maintenance, regulation of meristem growth. *VRS2* associated genes contained terms including regulation of cell size, phloem or xylem histogenesis, auxin efflux, plant-type sporogenesis, floral organ development, carbohydrate derivative catabolic process. *VRS3* associated genes was enriched for terms including response to high light intensity, seed germination, ethylene-activated signaling pathway, fruit development, seed development. *VRS4* associated genes contained terms including response to reactive oxygen species, lipid droplet, response to temperature stimulus, response to abscisic acid, response to glucose, response to hydrogen peroxide. The integrated results indicate the connections between *VRS1*-4 and the differences of the GO terms for their associated genes.

To validate the quality of our RNA-Seq dataset, we conducted RT-qPCR for *VRS2* and *VRS4* at the AP and TP stages. As determined by RT-qPCR, the expression of *VRS2* during the AP stage was higher than at the TP stage, while *VRS4* expression was lower at the at AP stage relative to the TP stage (Fig. S16), in agreement with our RNA-Seq analysis.

### The regulation of spikelet fertility by *VRS1-5* genes

#### (1) Determination of spikelet fertility

The indeterminate nature of barley spikes may allow over 30 rachis nodes of spikelet primordia to form within one spike. The number of fertile rachis nodes per spike (grain setting) at physiological maturity is a key factor determining final grain yield in barley. However, before the final grain number is set at physiological maturity, spikelet primordia structures undergo a sophisticated development and abortion process. After reaching the maximum number of floret primordia, representing its yield potential, barley spikes initiate a floral degradation process that will determine the number of fertile florets before the tipping stage when pollination has already occurred. Following this degradation of spikelet primordia before anthesis, another one to three additional spikelets are usually lost after pollination until final grain number is reached at physiological maturity (PM). Maximum floret primordia number and fertile rachis node (the nodes with at least one grain) are two crucial points during this spikelet developmental process.

A GO enrichment analysis (Fig. 4b, Fig. S7, S8, S9, S10, Table S10) indicated that phytohormones were involved in spikelet development and degeneration in barley. Cytokinins has been strongly implicated in many aspects of development, ranging from grain number and size by regulating cell division and differentiation, shoot and root growth, apical dominance, senescence, fruit and seed development, to responses to biotic and abiotic stressors (13–16). Therefore, we exploited mutations in each of the VRS genes to assess the interaction between *VRS1-5* and cytokinins as well as their effects on spikelet fertility.

#### (2) Spikelet fertility phenotypes in *vrs1-5* mutants

The detailed information of *vrs1-5* mutants can be found in previous work (17). We determined maximum number of spikelet primordia node at the AP stage. We discovered that loss-of-function mutations in any single *VRS1-5* gene produced fewer spikelet primordia node per spike (*vrs1*, 31.8; *vrs2*, 31. 7; *vrs3*, 31.0; *vrs4*, 31.0; *vrs5*, 30.5) than the Bowman control (34.5), suggesting that *vrs* 1-5 mutants suppress spikelet primordia initiation in barley (Fig. 6b).

**Fig. 6.**
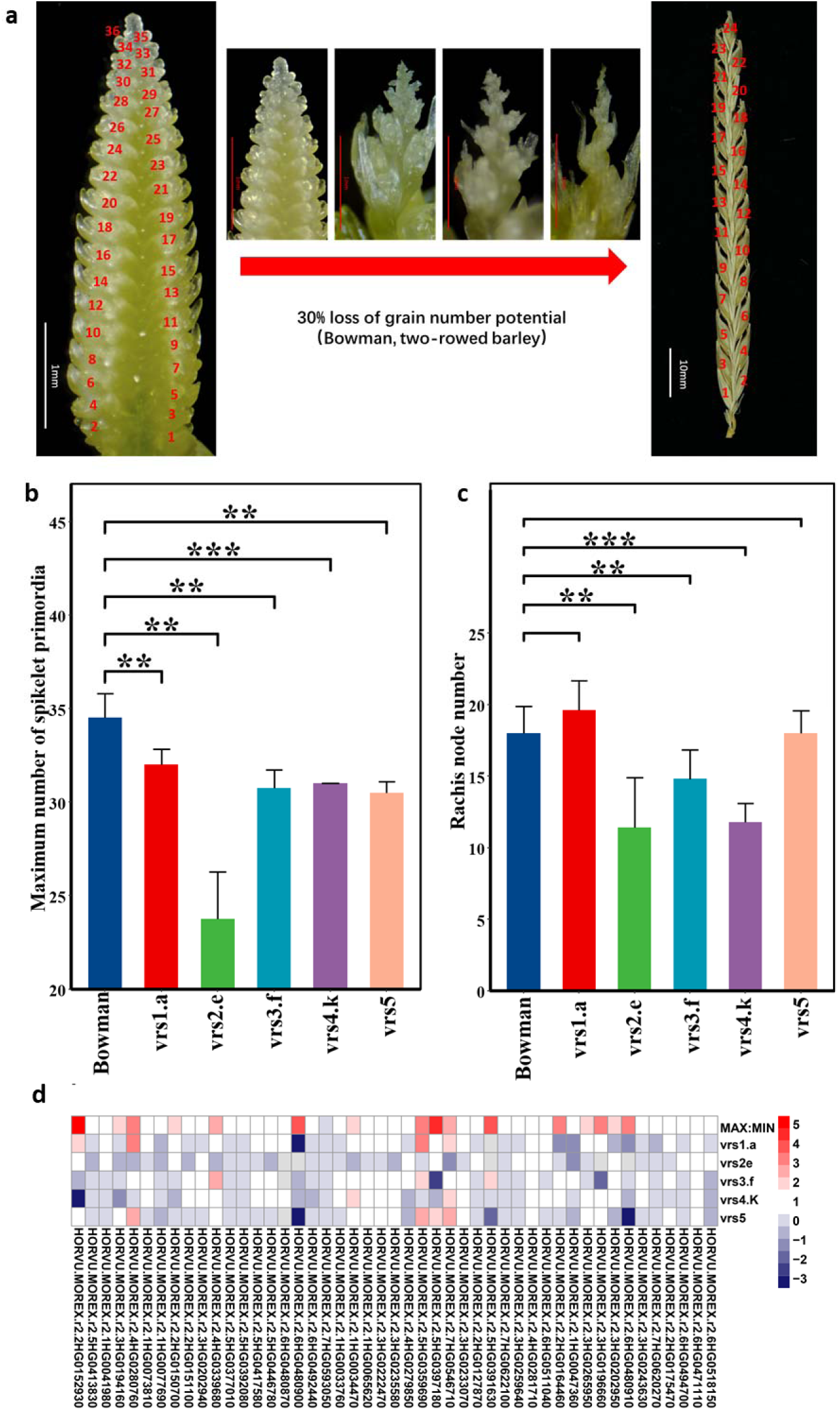
The effects of *VRS1-5* on spikelet fertility. (a) Loss of grain number potential. For Bowman (a two-rowed barley cultivar), the number of spikelet primordia reaches its maximum at the awn primordia stage (36 spikelet primordia per spike) and 24 spikelet primordia set grains at physiological maturity. (b, c) Differences in maximum number of spikelet primordia (b) and fertile rachis node number (c) between Bowman (control), *vrs1*.*a, vrs2*.*e, vrs3*.*f, vrs4*.*k* and *vrs5* genotypes. (d) Differences in gene expression for cytokinin metabolism and signaling between Bowman (control), *vrs1*.*a, vrs2*.*e, vrs3*.*f, vrs4*.*k* and *vrs 5* genotypes. In Fig. 6b, 6c, **, *** indicate significant differences (P-values of 0.01, 0.001, respectively). In Fig. 6d, the colors indicate the fold-change of gene expression induced by mutants.

Relative to the control (18.0 fertile rachis node per spike), vrs2, vrs3, vrs4 mutants lowered the number of fertile rachis node per spike (*vrs2*, 11.1; *vrs3*, 14.8; *vrs4*, 11.8), although we did not observe this phenotype for the *vrs1* (19.6) and *vrs5* (18.0) (Fig. 6c). Correspondingly, the survival rate of rachis nodes (the ratio between the maximum number of spikelet primordia node and the number of fertile spikelet nodes) was lower in *vrs2, vrs3*, and *vrs4* mutants (35%, 48%, 38%) than in the control (52%). These results indicate that *VRS2, VRS3* and *VRS4* play important roles in the determination of fertile rachis node number, whereas *VRS1* and *VRS5* contribute little to this trait.

#### (3) Phytohormones measurements in the wild type and *vrs* mutants

In addition to the effects on the number of rachis nodes per spike, we observed higher cytokinin levels in *vrs1-5* mutants (higher IP in *vrs1* mutant, 0.77; *vrs3* mutant, 0.82; *VRS4* mutant, 0.69; *vrs5* mutant, 0.89 and higher cZ in *vrs2* mutant, 0.06; relative to the control (IP 0.48, cZ, 0.19) (Fig. 17a), Indeed, *vrs1, vrs3, vrs4, vrs5* mutations increased cytokinin contents by 60%, 69%, 43% and 84%, respectively. We hypothesize that the higher cytokinin content resulting from the *vrs* mutations are responsible for their effects on spikelet node number.

#### (4) Phytohormones treatment in the wild type and *vrs* mutants

To investigate the effects of cytokinin on the number of rachis nodes, we treated the Bowman control and the *vrs1, vrs3, vrs4, vrs5* mutants with 6-Benzylaminopurine (6-BA) every 1–2 days starting at the AP stage. We determined that 6BA treatments lowered the number of fertile rachis nodes per spike (nodes with grains) in the *vrs1, vrs3, vrs4*, and *vrs5* mutants by 16%, 24%, 10%, and 30%, respectively. We did not observe a similar effect for the Bowman control (18.0 fertile spikelet nodes, without 6-BA treatment; 18.4 fertile spikelet nodes, with 6-BA treatment) (Fig. 17b). These results suggest that loss-of-function mutations in *VRS1, VRS2, VRS3, VRS4*, and *VRS5* increase the sensitivity of spikelet abortion to cytokinin treatment.

#### (5) RNA-Seq in the wild type and vrs mutants

To further investigate the connections between VRS and cytokinins, we conducted an RNA-Seq analysis of spikes at the YA stage, which corresponds to the pollination phase. We extracted the expression of 47 genes related to cytokinin biosynthesis and signaling according to the most recent released of the barley genome. We noticed that *vrs1-5* mutants greatly affected the expression of these 47 genes to various levels (Fig. 6d). This observation is consistent with our results above regarding endogenous cytokinin content and exogenous cytokinin treatment for spike number in this study.

## Discussion

The primary aims of this study were to explore the potential of VRS genes in raising barley grain yield. We first investigated possible correlations between haplotypes at each VRS and spike morphology traits in 894 worldwide barley accessions from 35 countries. We further conducted a high-resolution spatio-temporal transcriptome analysis of spikelet tissue over 17 stages and five positions, which allowed us to characterize the dynamic expression patterns underlying spike development and generate a co-expression network linking gene modules and spike morphology traits. Our phenotypic measurements, combined with our determination of the effects of cytokinin on wild-type and *vrs* mutant spikes provides foundational information for an investigation of the mechanism of spikelet primordia abortion.

Barley has been the target of intense artificial and natural selection since its domestication, resulting in large haplotypes, as observed in elite germplasm (18–21). This issue was also manifest in our work and illustrated by the *VRS1-5* haplotypes we observed across our barley collection of accessions. We detected these *VRS1-5* haplotypes in almost all countries represented by our collection, suggesting that these haplotypes have been widely used in barley breeding to increase yield and to broaden adaptation of variable environments worldwide.

Previous investigations have showed that *VRS1-5* mutants influence grain size and number (3, 17, 22). Therefore, knowing when and where *VRS1-5* are expressed during will provide vital information about their contribution to various spike development stages, especially during grain size enlargement and spikelet primordia degeneration, which normally results in a 30–50% loss of grain number potential. In the present work, we conducted a spatio-temporal transcriptome analysis to dissect the roles of *VRS1-5* on spike development. The variable *VRS1-5* expression patterns observed here suggest that they work during specific and distinct time windows of spikelet development and abortion after the AP stage. Previous studies had already described the expression patterns of *VRS1-5*(17, 23), but little information about their actions for spikelet fertility after the AP stage has been reported. In addition to *VRS1-5*, we characterized the expression patterns of other genes, which are consistent with the growth trends of spike components as determined by their associated enriched GO terms. The transcriptome dataset described here covered key developmental stages, from the AP stage to physiological maturity stage, and allowed us. to monitor carpel development and the grain filling process as well as spikelet primordia degeneration.

We analyzed *vrs1-5* mutants to dissect the effects associated with loss-of-function mutations in *VRS1-5* on the number of spikelet primordia nodes and fertile rachis nodes. Previous work had shown that inactivating *VRS1-5* may decrease the number of rachis node in barley (17). In agreement, we observed here fewer rachis nodes in *vrs1-5* mutants relative to the wild type. In addition, we noticed that *vrs1-5* mutations suppressed the initiation of apical spikelet primordia and accelerated the degeneration of spikelet primordia. Previous work has suggested that *VRS2, VRS3, VRS4* regulates the expression of genes involved in phytohormone metabolism, including that of cytokinins (3–5). In this study, cytokinin treatments on the wild type and mutants suggested that *VRS1-5* do regulate rachis node number by increasing cytokinin content in spikes and improving spikelet primordia sensitivity to cytokinins.

In summary, our results show that *VRS1-5* play independent and essential roles during spikelet development and, repression of apical spikelet primordia initiation. *VRS1-5* not only determine the spike row type, but can also be used to regulate spike components, which may further raise grain yield in barley. These results suggest that a better understanding of the processes that control spikelet development and degeneration may contribute to improving crop yield towards global food security.

## Methods and materials

### Growing condition and measurements of spike morphology traits for the resequencing barley populations

In this study, 894 spring barley varieties were selected for evaluating agronomic traits. The 894 barley accessions are originally from the 35 countries worldwide (Table S1).

We planted the plants in four conditions: three field conditions and one greenhouse condition. Field experiments were carried out during 2019-2020 growing season at three locations: Beijing (116.30°E, 39.90°N), Yancheng (120.54°E, 34.28°N) and Chongzhou (103.67°E, 30.63°N). Around 200 plants per accession (for all the 894 accessions) were planted in these three field conditions. For the greenhouse experiment, we selected 390 accessions and 20 plants for each accession were planted for the measurements of spike morphology traits.

For the greenhouse experiment, we germinated the seeds in petri dish (photoperiod, 16 h/8 h, light/dark; temperature, 20°C/16°C, light/dark) for four days. We then vernalized germinated seeds at 4 °C for 28 days. After that, we transplanted four seedlings to one pot (19×19×18cm, 6.5 liters). Supplemental light was supplied and plants were irrigated when required. The spike length (cm), spike chaff per spike (g), rachis dry weight per spike (g), awn dry weight per spike (g), spike DW (g), grain weight per spike (g), grain number per spike, and total and fertile rachis node numbers were measured at physiological maturity. Six plants for each cultivar were randomly selected for trait measurements. The spike length was measured without the awn. The rachis dry weight per spike refers to the rachis after removal of spikelets. The density of rachis node was determined as the ratio between rachis node number per spike and spike length. In addition, we calculated the ratio between awn dry weight and spike dry weight, the ratio between grain weight and rachis dry weight, the ratio between spike dry weight and rachis dry weight, the ratio between grain dry weight and awn dry weight as well as the ratio between awn dry weight and spike rachis dry weight in this study.

### Growing condition and sample collections for transcriptome analysis

Six-rowed barley (*Hordeum vulgare* cv Morex) plants were grown in a greenhouse. We germinated the seeds in petri dish (photoperiod, 16 h/8 h, light/dark; temperature, 20°C/16°C, light/dark) for four days. We then vernalized germinated seeds at 4 °C for 28 days. After that, we transplanted four seedlings to one pot (19×19×18cm, 6.5 liters). Supplemental light was supplied and plants were irrigated when required.

We conducted spatiotemporal dissection to reveal the overall information of floret development and abortion at 17 spikelet developmental stages in barley: awn primordium (AP) stage (spikelet primordium number reach maximum, the apical meristematic dome stops activity and no more primordia will initiate), white anther (WA) stage (the anthers partially surround by the young carpel, the two bumps of top carpel initiate the development of styles and stigma, the palea particallly encloses the carpel and stamen, the rachilla can be seed between palea and lemma), green anther (GA) stage (the stamens grow to around 1mm long, the carpel is enclosed by the lemma and palea). Yellow anther (YA) stage (the anthers and pollens are more advanced, the lemma and palea grow and completely enclose the stamens and carpel), tipping (TP) stage (the tips of awns are visible), early heading (EHD) stage (The tip of top spikelet visible), middle heading (MHD) stage (the half of spike visible) late heading (LHD) stages (more that 90% of spike visible). The information to determine the different developmental stages can refer to (24, 25). We took all the three spikelets at each rachis node at the 17 stages for the temporal transcriptome analysis of spikelet (Fig. 1). We took all the three spikelets at each rachis node at five positions of rachis node ((0-10%, 30%, 50%, 70%, and 90-100% of spike length, Fig. 1) within individual spike for the spatial transcriptome analysis of spikelet (Fig. 1). Three biological replicates were set up for each of the time/space points. Each replicate was obtained by pooling samples from at least five to ten plants. Total RNA was extracted using TRIzol reagent (Invitrogen).

At AP stage, main shoots of three plants were randomly selected to determine maximum number of floret primordium per spike. At physiological maturity, main shoots of three plants were randomly selected to determine rachis node number per spike. At all the 17 spike developmental stages, we measured spike length and spike dry weight. At the 14 stages (without AP, GA, YA stages) and five positions, we measured awn dry weight, spikelet dry weight (without awn), central spikelet dry weight (without awn), lateral spikelet dry weight (without awn). These traits are too small for the dry weight measurement at AP, GA and YA stage, therefore we did not determine these traits at the three stages.

### PCR amplification, sequencing and haplotype analysis of *VRS1-5* genes

DNA extraction, quantification and qualification were conducted as previously described (26). In brief, 100-200 mg of the seedling leaves in the field (Nanyang Research Station, Yancheng, 2018) were sampled for DNA miniprep using a modified Cetyltrimethyl Ammonium Bromide (CTAB)-based DNA isolation approach (27). DNA quality and concentration were measured using a NanoDrop spectrophotometer (Thermo Scientific, Wilmington, USA) and gel electrophoresis by using λDNA. The original DNA stocks were stored at -20°C, while a few aliquots were diluted to the working concentration (10-20 ng/μl) for PCR reaction. Sequences from each accession were amplified from genomic DNA (gDNA). Primer pairs used to amplify the longest exon sequences of *VRS1-5* were designed using Primer-Blast in NCBI (Table S12). PCR amplification in a reaction volume of 12.5 μl, containing 40 ng template DNA, 1 × Phanta Max Buffer and 0.1 U Phanta Max Super-Fidelity DNA polymerase (Mg^2+^ plus) (Vazyme, Nanjing, China), 0.2mM dNTPs and 0.4 μM of each primer. The PCR procedure started for initial denaturation at 95° C for 5 min, following 36 cycles including 15 s for denaturation at 95° C 20 s for annealing at 60° C and 1 min for extension at 72 ° C (extension rate is 1 kb per minute), followed by a final extension at 72 °C for 5 min. Ten microliter of each PCR amplicon was sequenced and the remaining was separated for specificity test by gel electrophoresis on 1% agarose gels (160 voltage). To obtain SNP for the 894 barley accession, multiple sequence alignment was conducted in DNAMAN Version7 and TASSEL5.0 software was further used to get SNPs (28). DnaSP6 was further used to determine the number of haplotypes, haplotype diversity, nucleotide diversity. Analysis of variance (ANOVA) for the spike morphology traits in the four environments was conducted in sommer package in R (29).

### RNA-Seq

The spikes at YA stage were frozen in liquid nitrogen, ground into powder, total RNA was extracted with TRIzol Reagent (Invitrogen), treated with DNase and purified with RNeasy columns (QIAGEN). A total amount of 1 µg RNA per sample was used as input material for the RNA sample preparations. Sequencing libraries were generated using NEBNext UltraTM RNA Library Prep Kit for Illumina (NEB, USA) following manufacturer’s recommendations and index codes were added to attribute sequences to each sample. The clustering of the index-coded samples was performed on a cBot Cluster Generation System using TruSeq PE Cluster Kit v3-cBot-HS (Illumia) according to the manufacturer’s instructions. After cluster generation, the library preparations were sequenced on an Illumina Novaseq platform and 150 bp paired-end reads were generated.

### Read mapping and transcript profiling

The barley reference genome was downloaded from https://doi.ipk-gatersleben.de/DOI/83e8e186-dc4b-47f7-a820-28ad37cb176b/64687610-b20c-4702-9698-7dc2401a80e5/0. The quality of raw data was assessed using Fastqc (http://www.bioinformatics.babraham.ac.uk/projects/fastqc/), after that, raw RNA-Seq reads were processed to remove low-quality reads using trimmomatic (Trimmomatic: a flexible trimmer for Illumina sequence data). Then we used salmon (30) and the tximport package in R (31) to quantify RNA-seq data and import salmon’s transcript-level quantifications and optionally aggregate them to the gene level for differential expression analysis. We further used DESeq2 in R (32) with the optional fpkm() to obtain Fragments Per Kilobase of exon model per Million mapped fragments (FPKM) for each sample. ImpulseDE2 (v1.6.1) in R (33) was used to identify significant dynamic genes from AP stage to 36DFL stage at each spikelet position. ImpulseDE2 was used in case-only mode and no batch effects settings were added to the parameters.

### Gene annotation and functional enrichment analysis

To obtain gene descriptions, we used precomputed clusters and phylogenies from the eggNOG database for the functional annotation of obtained sequences though a tool called eggNOG-mapper. Gene Ontology (GO) enrichment analyses were performed using clusterProfile package in R (34).

### Module-trait association analyses

Module-trait association analyses were performed using the WGCNA package (v1.51) in R (35). The top 75% of the median absolute deviation (MAD) were screened, remaining 30060 genes were used for the analysis. The module-trait associations were obtained using a one-step network construction method construction function (blockwiseModules) with default settings, except that the soft threshold (power) was 17, TOMType was unsigned, minModuleSize was 34, and mergeCutHeight was 0.25. The relationships between modules and traits were assessed using the correlations between the module eigengene and the trait, allowing the identification of modules that are correlated with traits and searching for the most significant associations.

### qRT-PCR Expression Analysis of Genes

Total RNA was extracted with Omega plant RNA kit (Code No. R6827-01). Reverse transcription was performed with 2 μg RNA using ReverTra Ace qPCR RT Kit (Code No. FSQ-101). Real-time PCR was performed using SYBR Premix Ex Taq II (Takara Bio) on an ABI 7500 Real-Time PCR system (Applied Biosystems). Barly ACTIN was used as an endogenous reference gene to normalize the data. Three technical and three biological replicates were used to produce the average expression levels relative to the reference. The primers of *VRS2* and *VRS4* can be found in previous work (4, 5) (Table S13).

### Determination of endogenous Cytokinin concentration

The spikes at YA stage were frozen in liquid nitrogen, ground into powder, and extracted with methanol/water/formic acid (15:4:1, V/V/V). The combined extracts were evaporated to dryness under a nitrogen gas stream, reconstituted in 80% methanol (V/V), and filtered (PTFE, 0.22 μm; Anpel). The sample extracts were analyzed using an LC-ESI-MS/MS system (HPLC, Shim-pack UFLC SHIMADZU CBM30A system; MS, Applied Biosystems 6500 Triple), and cytokinin content were detected by MetWare (http://www.metware.cn/) based on the AB Sciex QTRAP 6500 LC-MS/MS platform. Three replicates of each assay were performed.

## Acknowledgements

We thank Sarah M. McKim (The James Hutton Institute) for providing seeds of mutants. We thank Leibniz Institute of Plant Genetics and Crop Plant Research for providing greenhouse space for the collections of transcriptome samples. This work was supported by a grant to Z.G., L.Z. and L.S. from Key Laboratory of Plant Molecular Physiology, Institute of Botany, Chinese Academy of Sciences. P.Y. is financed by Fundamental Research Funds for Central Non-Profit of Institute of Crop Sciences of CAAS and Agricultural Science and Technology Innovation Program of CAAS.

## Author contributions

Z.G. conceived the project. Z.G. and P.Y. designed and supervised the experiments. Y.L., Z.W., Z.S., L.Z., H.W., K.Y. and L.S. measured the spike morphology traits. L.S. conducted the exogenous hormones treatments and qRT-PCR analysis. Y.C. carried out DNA isolation as well as plant sowing, management and sampling in the field, Z.G. wrote the manuscript. Y.L. and Y.J. performed data analyses. All the authors viewed and edited the manuscript.

## Competing interests

The authors declare no competing interests.

## Notes

### Competing Interest Statement

The authors have declared no competing interest.

